# Strategic coexistence theory for evolutionary games

**DOI:** 10.64898/2026.06.24.734261

**Authors:** Sang Woo Park

## Abstract

Evolutionary game theory and ecological coexistence theory both seek to predict the outcome of competition between biological entities, be they strategies or species, but the two fields have relied on largely separate approaches. Replicator equations provide a foundation for analyzing strategy competition, yet they do not explicitly separate the mechanisms that stabilize competition from those that equalize fitness differences between strategies. Here, we extend modern coexistence theory from community ecology to develop strategic coexistence theory (SCT), a framework for quantifying strategic niche and fitness differences between competing strategies. SCT recovers the classic classification of two-strategy games, distinguishing competitive exclusion, coexistence, and priority effects within a shared niche–fitness difference space. Applying SCT to five mechanisms for the evolution of cooperation further reveals that these mechanisms promote cooperation through distinct dynamical routes: kin selection, network reciprocity, and group selection primarily reduce fitness differences, whereas direct and indirect reciprocity destabilize competition and generate priority effects. Finally, applying SCT to microbial public-goods game shows that nonlinear microbial growth can both stabilize and equalize competition between co-operators and defectors, allowing coexistence. Together, these results show that SCT provides a complementary framework for comparing evolutionary games and teasing apart the coexistence mechanisms underlying strategy competition.

## Introduction

Understanding the mechanisms that promote the evolutionary stability of competing strategies is a fundamental aim in evolutionary biology (Lewontin, 1961; Slobodkin, 1964; Smith and Price, 1973; Taylor and Jonker, 1978). To this end, evolutionary game theory (EGT) and replicator dynamics have provided key foundations, allowing us to predict the outcome of strategic competition through payoff inequalities, invasion criteria, and stability analysis (Schuster and Sigmund, 1983; Hofbauer and Sigmund, 1998; Cressman, 2003; Cressman and Tao, 2014). However, less attention has been given to similarities between replicator dynamics and community ecology until recently (Gokhale and Traulsen, 2025; Tarnita and Traulsen, 2025; Allesina, 2026), even though the classic replicator equations and the generalized Lotka-Volterra equation are mathematically equivalent (Hofbauer, 1981; Page and Nowak, 2002).

Interestingly, even a simple two-strategy game can yield heterogeneous outcomes that mirror outcomes in community ecology (Hauert, 2002; Szabó and Fath, 2007; Cooney, 2020; Tarnita and Traulsen, 2025): the dominance of a single strategy (prisoner’s dilemma and harmony games), coexistence (hawk–dove games), and bistability (stag-hunt games). Such differences in conditions that promote evolutionary stability and dominance of a single strategy and those that permit strategy coexistence indicate that EGT can be understood from the perspective of coexistence theory (Motro, 1991; Doebeli et al., 2004; Hauert and Doebeli, 2004; Doebeli and Hauert, 2005; Gore et al., 2009; Souza et al., 2009; Archetti and Scheuring, 2011; Steidinger and Bever, 2014; Tarnita and Traulsen, 2025). In fact, stable polymorphisms of evolutionary strategies, such as coexistence of cooperators and defectors, are observed in microbial and other biological systems (Gore et al., 2009; Cordero et al., 2012; Kocher et al., 2018; Pollak et al., 2021; Lin et al., 2024), but coexistence-theoretic perspective has often remained implicit. Recasting game dynamics as a coexistence problem immediately raises the following question: when two strategies coexist, does the corresponding mechanism stabilize or equalize competition (Chesson, 2000)? While EGT allows us to predict whether two strategies can coexist or not, a framework for quantifying coexistence mechanisms between competing strategies remains elusive.

Modern coexistence theory (MCT) from community ecology presents a particularly useful framework for quantifying coexistence mechanisms (Chesson, 2000; Adler et al., 2007; Godoy et al., 2014; Godoy and Levine, 2014; Kraft et al., 2015; Adler et al., 2026). MCT predicts that coexistence of two competing entities, be they species or strategies, requires stabilizing niche differences to overcome average fitness differences (Chesson, 2000; Adler et al., 2007; Kraft et al., 2015). In community ecology, stabilizing niche differences represent differences in factors that limit the growth of competing species, causing each species to limit itself more than it limits its competitor (Chesson, 2000). In EGT, a strategy’s niche can be interpreted as strategic environments defined by the composition of competing strategies and the resulting interactions (Taylor and Jonker, 1978; Cressman and Tao, 2014; Tarnita and Traulsen, 2025). Thus, stabilizing niche differences arise when each strategy is favored by a different strategic environment, such that the spread of each strategy creates conditions that limit its own relative growth and favor the recovery of its competitor. In contrast, fitness differences represent differences in the ability of each entity to compete under the limiting environments created by its competitor (Chesson, 2000). This differs from payoff differences in EGT, which depend on the underlying strategy frequency and therefore combine both stabilizing effects caused by changes in the strategic environment and residual fitness differences that favor one strategy over another (Taylor and Jonker, 1978; Cressman and Tao, 2014; Tarnita and Traulsen, 2025).

So far, MCT has been primarily utilized for understanding species coexistence, but recent work showed that MCT can be extended beyond species competition, especially for studying pathogen strain competition, allowing one to directly quantify niche and fitness differences of competing entities, be they species or strains (Sieben et al., 2022; Park et al., 2024, 2026). Inspired by this effort, we present a Strategic Coexistence Theory (SCT) for predicting coexistence and exclusion of competing strategies in EGT by quantifying strategic niche and fitness differences. We begin by introducing SCT, which can be applied to any generic system of EGT described by replicator equations. We show that SCT can recover outcomes from classic, two-strategy games, providing a reinterpretation of classic results from coexistence-theoretic perspective. We then apply SCT to comparing five mechanisms for the evolution of cooperation, revealing that direct and indirect reciprocity destabilize competition between cooperation and defection, thus pushing the system towards priority effect regime. Finally, we consider a case study of microbial cooperation, where nonlinear microbial growth can both stabilize and equalize competition between cooperation and defection, allowing coexistence of strategies.

### Strategic coexistence theory

Here, we present a brief derivation of strategic coexistence theory (SCT) (Figure 1). To do so, we begin with a classic replicator equation, which allows us to model changes in the frequency of competing strategies based on their payoffs (Schuster and Sigmund, 1983; Hofbauer and Sigmund, 1998; Cressman, 2003; Cressman and Tao, 2014):

**Figure 1.**
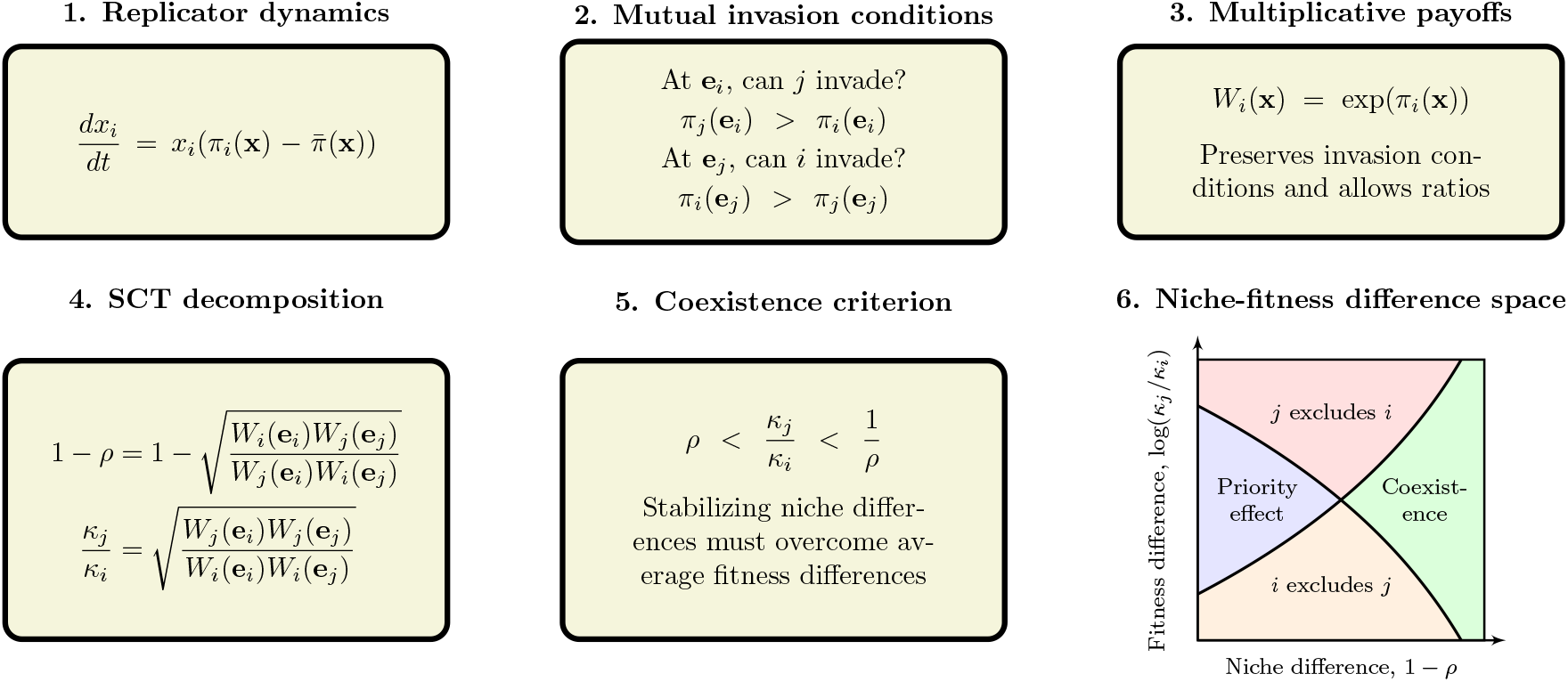
A schematic diagram summarizing strategic coexistence theory. Strategic coexistence theory decomposes pairwise replicator dynamics into stabilizing and equalizing mechanisms. Mutual invasion conditions can be rewritten on a multiplicative payoff scale, allowing us to derive strategic niche difference 1 − *ρ* and fitness difference *κ*_*j*_*/κ*_*i*_ based on multiplicative payoff ratios. Then, the inequality *ρ < κ*_*j*_*/κ*_*i*_ *<* 1*/ρ* determines mutual invasion.

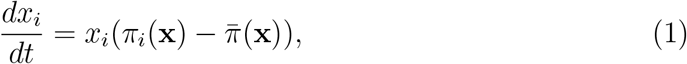

where *x*_*i*_ represents the frequency of strategy *i, π*_*i*_(**x**) represents the payoff of strategy *i* for a given strategy distribution **x**, and 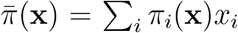 represents the average payoff. Since strategy frequencies must sum to 1 (i.e.,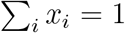), a single-strategy equilibrium dominated by strategy *i* can be described by a unit vector **e**_*i*_, where all elements are equal to zero except for the *i*-th element. We use these single-strategy equilibria to define niche and fitness differences between two competing strategies *i* and *j*.

Because payoffs in many games can be negative, payoff ratios can be difficult to interpret. Thus, we work on a positive multiplicative scale by exponentiating the payoff terms:

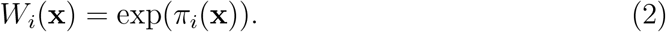

This exponential transformation is convenient because it preserves the payoff ordering and does not alter the invasion conditions. Therefore, this allows us to compare strategic environments through ratios, as in modern coexistence theory. For convenience, we refer to *W*_*i*_(**x**) as the multiplicative payoff of strategy *i*.

First, we define the niche overlap *ρ* between strategies *i* and *j* as the ratio between the geometric mean of each strategy’s multiplicative payoff in the strategic environment that it creates when common, 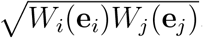, and the geometric mean of each strategy’s multiplicative payoff in the environment generated by its competitor, 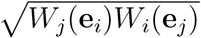. The corresponding niche difference is then given by

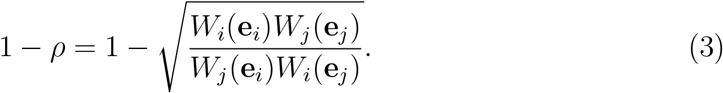

When 1 − *ρ >* 0, each strategy performs better, on average, in the environment generated by its competitor than in the environment generated by itself. In other words, intraspecific competition is stronger than interspecific competition. Second, we define the fitness difference as the ratio between the geometric means of the multiplicative payoff of each strategy across the two single-strategy environments:

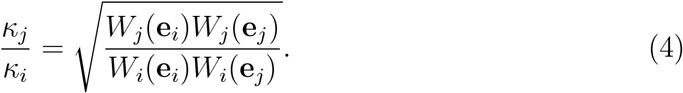

In other words, strategy *j* has an average competitive advantage over strategy *i* (*κ*_*j*_*/κ*_*i*_ *>* 1) when it performs better, on average, across the environments generated by itself and its competitor.

Together, for any two strategies described by the classic replicator equation, mutual invasion requires stabilizing niche differences to overcome average fitness differences. Specifically, the condition for mutual invasion can be rewritten as (Chesson, 2000):

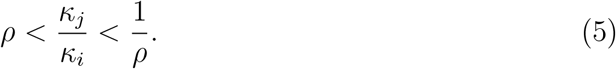

This inequality applies to any games described by the classic replicator equation because *ρ* and *κ*_*j*_*/κ*_*i*_ can be calculated directly from ratios of multiplicative payoffs. This decomposition not only allows us to understand how different mechanisms contribute to the coexistence or exclusion of competing strategies but also provides shared axes for comparing different games. Detailed derivations are provided in Supplementary Text S1.

We note that the absolute magnitudes of niche and fitness differences are sensitive to payoff scaling because they are defined on an exponentiated payoff scale. However, the qualitative outcome of strategic competition under SCT is preserved because the exponential transformation does not alter invasion conditions.

### Classic two-strategy games

As an illustrative example, we begin with classic two-strategy games (Hauert, 2002; Szabó and Fath, 2007; Cooney, 2020; Tarnita and Traulsen, 2025), which can be described by the following payoff matrix (Figure 2A):

**Figure 2.**
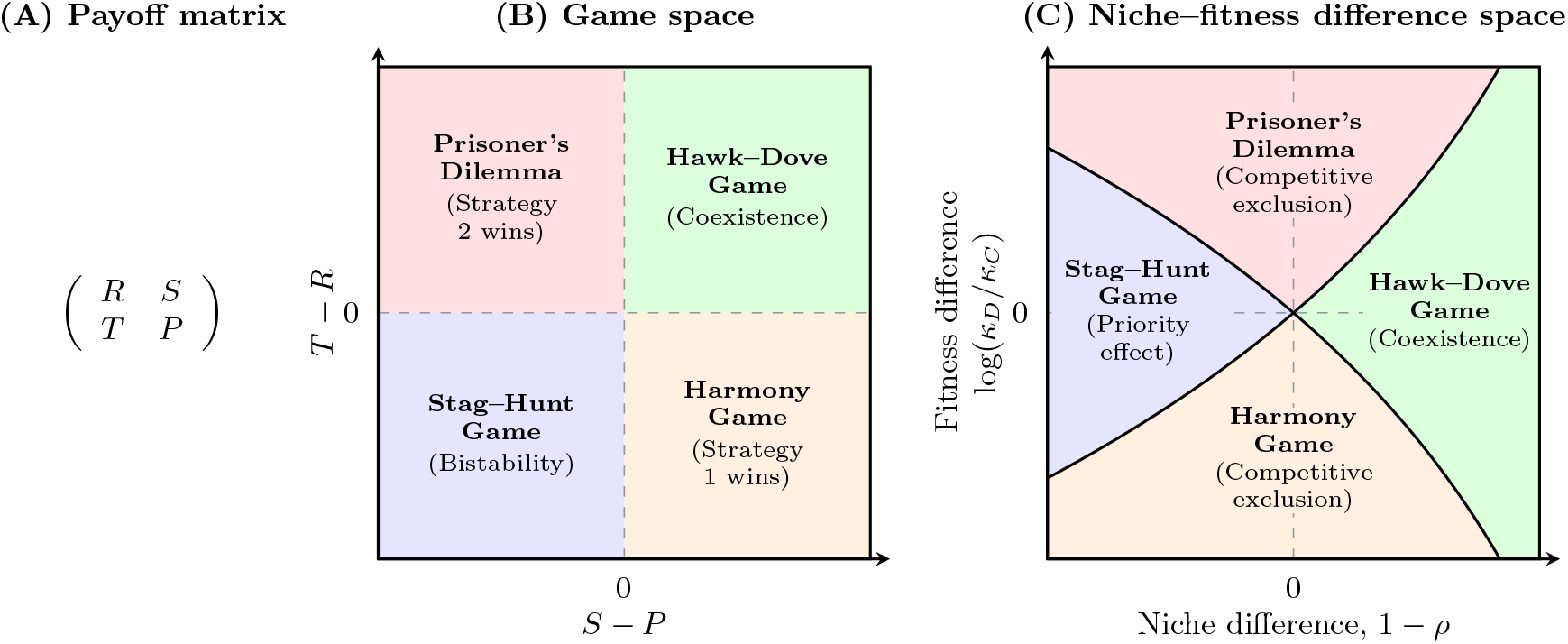
Coexistence-theoretic interpretation of classic two-strategy games. (A) General payoff matrix for a two-strategy game, where rows represent the focal strategy and columns represent the strategy of the interacting opponent. (B) Classification of four classic games based on the payoff differences *T* − *R* and *S* − *P* . Prisoner’s dilemma and harmony games lead to the dominance of one strategy; hawk–dove games allow coexistence; and stag–hunt games result in bistability. (C) Mapping of the same four games onto niche–fitness difference space. Hawk– dove games correspond to the coexistence regime, where stabilizing niche differences overcome fitness differences. Prisoner’s dilemma and harmony games correspond to competitive-exclusion regimes, whereas stag–hunt games correspond to the priority-effect regime. *R > P* is assumed throughout, but this does not affect coexistence outcome in any way.

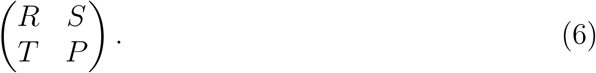

Here, the first and second rows represent strategies 1 and 2, respectively, and the columns represent the strategy of the interacting opponent. Writing *x*_1_ to denote the frequency of strategy 1, the expected payoffs are

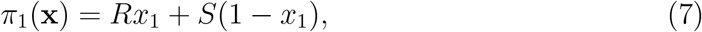

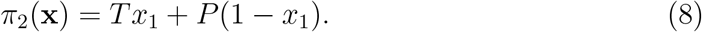

Thus, the dynamics can be written as

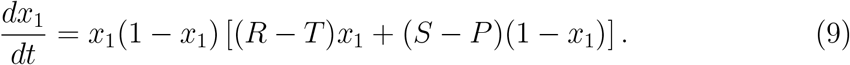

The signs of *T* − *R* and *S* − *P* determine whether each strategy can invade the other when rare. Specifically, strategy 2 can invade strategy 1 when *T > R*, whereas strategy 1 can invade strategy 2 when *S > P* . These two invasion conditions partition the game space into four regimes (Figure 2B): prisoner’s dilemma (*T > R > P > S*), hawk–dove game (*T > R, S > P* ), stag–hunt game (*R > T, P > S*), and harmony game (*R > T, S > P* ). In standard EGT, these outcomes are obtained by analyzing the stability of equilibrium conditions. For simple games, this analysis is straightforward.

A coexistence-theoretic approach recovers the same classification but provides a different interpretation (Figure 2C). For a two-strategy game determined by this payoff matrix, the niche difference and fitness difference are given by the following set of equations (Supplementary Text S2):

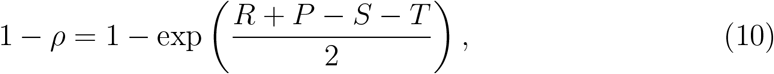

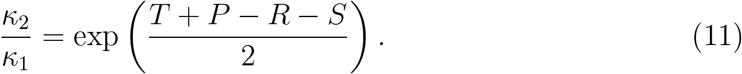

As shown above, mutual invasion requires *ρ < κ*_2_*/κ*_1_ *<* 1*/ρ*, which is equivalent to *T > R* and *S > P* . This corresponds to the hawk–dove game, in which each strategy can invade its competitor when rare and the two strategies coexist. The remaining games also have natural interpretations in niche–fitness difference space. Prisoner’s dilemma and harmony games represent competitive exclusion regimes, where one strategy has a fitness advantage that exceeds niche differences. In contrast, stag–hunt games represent a priority-effect regime, where neither strategy can invade the other and the outcome depends on initial conditions. Therefore, coexistence theory not only allows us to recover the classic results from two-strategy games but also provides new interpretations for these results in terms of niche and fitness differences. Building on this, we next apply this framework to comparing core mechanisms of cooperation.

### Comparisons of mechanisms for the evolution of co-operation

Even though cooperative behavior can be found across a wide range of biological systems, a simple prisoner’s dilemma shows that cooperation is not an evolutionarily stable strategy (Axelrod and Hamilton, 1981; May, 1987). Thus, understanding mechanisms that promote the evolution of cooperation has remained a central goal in evolutionary biology (West et al., 2007; Fletcher and Doebeli, 2008; Clutton-Brock, 2009; Kingma et al., 2014; Rakoff-Nahoum et al., 2016; Hilbe et al., 2018; Apicella and Silk, 2019; West et al., 2021; Lee et al., 2022; Palmer and Foster, 2022; Michel-Mata et al., 2024). Here, we compare five classic mechanisms for the evolution of cooperation using SCT (Nowak, 2006) and ask how they differ from the perspective of SCT.

Specifically, we consider the following mechanisms: kin selection, direct reciprocity, indirect reciprocity, network reciprocity, and group selection (Figure 3A). Kin selection favors cooperation when two interacting individuals are genetic relatives, where the genetic relatedness *r* must be greater than the cost-to-benefit ratio: *r > c/b* (Hamilton, 1964). Direct reciprocity favors cooperation when current co-operation may promote future cooperation by the same individual through repeated encounters (Trivers, 1971; Axelrod and Hamilton, 1981). In this case, the probability of repeated interaction (i.e., “next round”) must be greater than the cost-to-benefit ratio: *w > c/b*. Indirect reciprocity favors cooperation when current cooperation may promote future cooperation by different individuals through reputation (Nowak and Sigmund, 1998; Wedekind and Milinski, 2000; Nowak and Sigmund, 2005). Thus, the social acquaintanceship (i.e., the probability of knowing someone’s reputation) must be greater than the cost-to-benefit ratio: *q > c/b*. Network reciprocity favors cooperation via clustering of cooperators, requiring the benefit-to-cost ratio to be greater than the average number of neighbors: *b/c > k* (Nowak and May, 1992; Lieberman et al., 2005). Finally, group selection favors cooperation via multilevel selection, where groups with more cooperators exhibit faster growth at the group level even if defectors reproduce faster at the individual level (Traulsen and Nowak, 2006). This requires *b/c >* 1 + (*n/m*), where *n* is the maximum group size and *m* is the number of groups. All of these mechanisms can be described by a simple 2 × 2 payoff matrix (Nowak, 2006), from which the corresponding niche and fitness differences can be computed using Eq. (10) and Eq. (11). Details on niche and fitness difference calculations are presented in Supplementary Text S3.

**Figure 3.**
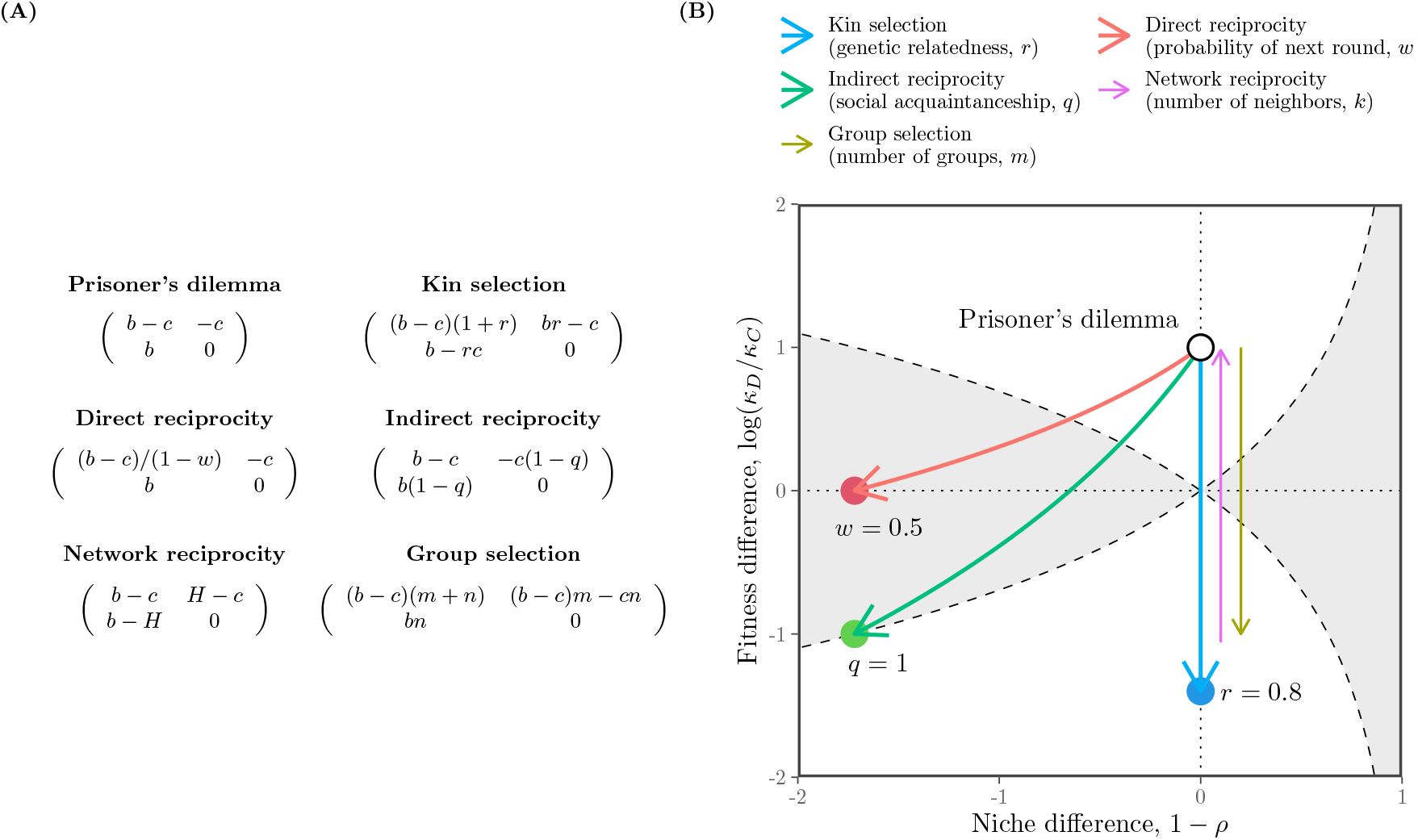
Comparisons of five mechanisms for the evolution of cooperation using SCT. (A) Payoff matrices for the baseline scenario (prisoner’s dilemma) and five mechanisms for the evolution of cooperation (Nowak, 2006). Prisoner’s dilemma is characterized by the benefit for the recipient *b* and the cost for the donor *c*. Network reciprocity is parameterized based on *H* = [(*b* − *c*)*k* − 2*c*]*/*[(*k* +1)(*k* − 2)]. Parameters for all other mechanisms are described in panel B. (B) Changes in niche and fitness differences driven by each mechanism. Each arrow represents the direction of change as we increase the corresponding parameter, indicated in figure legends. Note that kin selection, network reciprocity, and group selection all result in changes in fitness difference without causing any changes in niche difference. Therefore, we indicate the direction of change for network reciprocity and group selection separately to avoid overlapping arrows. For simplicity, we focus on the effect of number of groups *m* for group selection; increasing the number of maximum group size *n* has the opposite effect (Supplementary Text S3).

First, the prisoner’s dilemma provides a baseline scenario, where defection is evolutionarily stable (Figure 3A). From the perspective of SCT, there is complete niche overlap, and therefore defection competitively excludes cooperation (Figure 3B). Building on this, SCT shows that five mechanisms promote evolutionary stability of cooperation via different dynamical routes (Figure 3B). For example, kin selection allows the fitness of cooperation to increase as genetic relatedness *r* increases (Figure 3B). However, it does not affect the niche difference between cooperation and defection. Similarly, network reciprocity and group selection modulate the fitness of cooperation without causing any changes in strategic niches. In contrast, direct and indirect reciprocity destabilize competition while increasing the fitness of cooperation, thereby pushing the competition outcome towards a priority effect regime.

Equivalently, these two mechanisms promote bistability of defection and cooperation. SCT further reveals another difference between direct and indirect reciprocity. In particular, direct reciprocity is unable to generate sufficient fitness difference to competitively exclude defection even when the probability of next round *w* reaches 1 (Supplementary Text S3). In contrast, indirect reciprocity is able to generate sufficient fitness difference to push the system towards the boundary between priority-effect regime and competitive-exclusion regime as social acquaintanceship *q* reaches 1, indicating that an additional mechanism may be able to promote complete dominance of cooperation (Supplementary Text S3).

We note that none of these mechanisms allow coexistence of cooperation and defection. In many biological systems, such coexistence is often observed (Gore et al., 2009; Cordero et al., 2012; Kocher et al., 2018; Pollak et al., 2021; Lin et al., 2024). Therefore, we move on to the final example, applying SCT to microbial cooperation, where nonlinear benefits and environmental context can promote coexistence of cooperators and defectors.

### Microbial cooperation and public goods

Here, we consider a simple, phenomenological model of competition between cooperating and cheating yeast in a cooperative sucrose-metabolism system, where invertase production by cooperators allows cheaters to utilize glucose, a public good released through sucrose hydrolysis (Carlson and Botstein, 1982; Dickinson, 1998; Gore et al., 2009). Specifically, invertase production by cooperators incurs a cost *c* and generates a total benefit of 1 that is captured privately with efficiency *ϵ*. Assuming a linear relationship between glucose availability and microbial growth, the payoffs for cooperators *π*_*C*_ and cheaters (defectors) *π*_*D*_ can then be written as

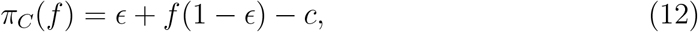

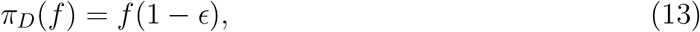

where the use of glucose depends on the fraction of cooperators *f* . The dynamics of cooperators and defectors are described by the following replicator equation:

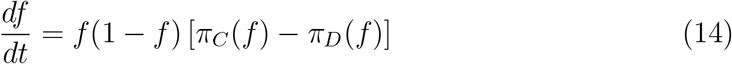

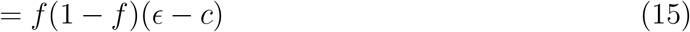

For this linear growth model, the payoff difference is fixed to *ϵ* − *c* independent of *f* (Figure 4A). Therefore, cooperation is favored when efficiency is greater than the cost of cooperation, *ϵ > c*, whereas defection is favored when the cost is greater than efficiency, *c > ϵ* (Gore et al., 2009).

**Figure 4.**
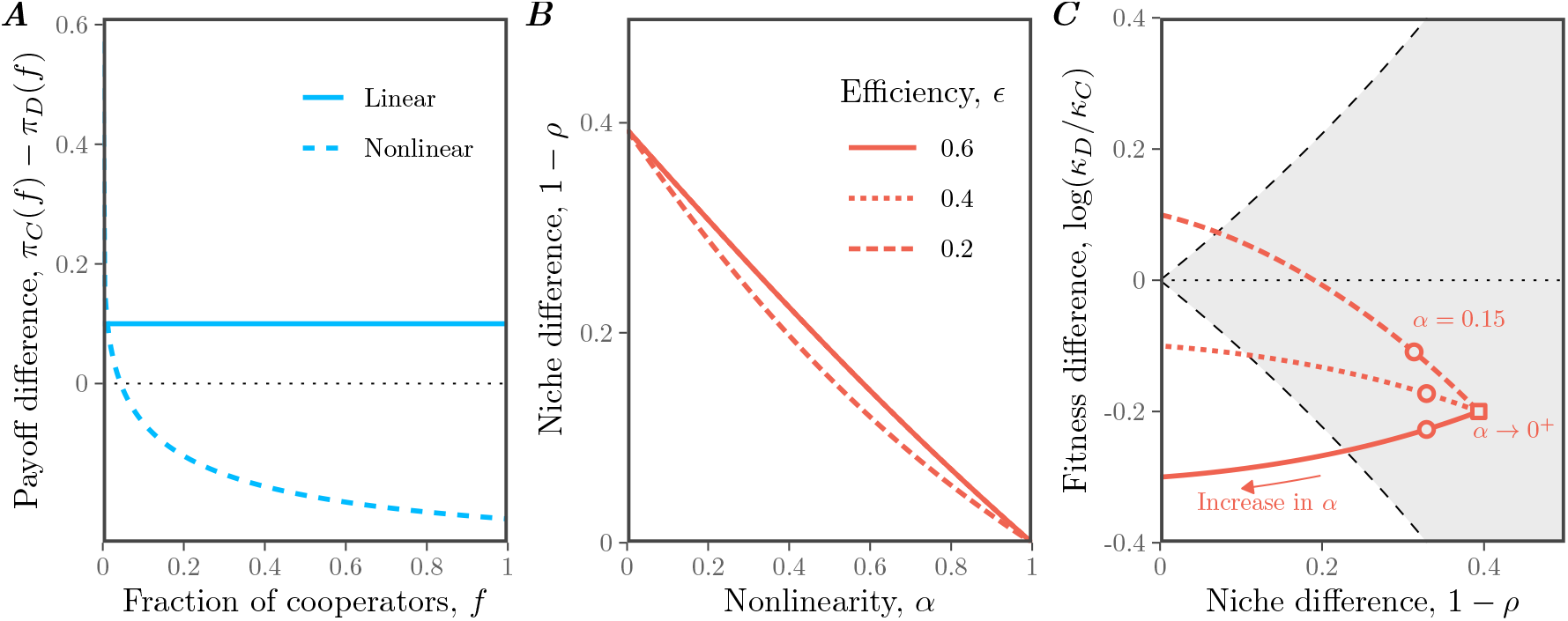
Effects of nonlinear microbial growth on coexistence of cooperation and defection. (A) Payoff differences between cooperation and defection for linear and nonlinear models of microbial growth. In panel A, we assumed *ϵ* = 0.4, *c* = 0.3, and *α* = 0.15 for illustrative purposes. (B) Effect of nonlinearity *α* on niche difference 1 *− ρ*. Each line assumes a different value of efficiency *ϵ*. Note that lines for *ϵ* = 0.6 and *ϵ* = 0.4 are nearly overlapping and are indistinguishable. (C) Effect of nonlinearity *α* on both niche difference 1 *− ρ* and fitness difference log(*κ*_*D*_*/κ*_*C*_). The gray shaded area represents the coexistence regime. The square represents the condition when *α* → 0^+^. The circles represent the condition when *α* = 0.15, which correspond to the experimental relationship between glucose and microbial growth (Gore et al., 2009). In panels B and C, we assumed *c* = 0.3, while allowing *ϵ* and *α* to vary.

Our framework allows us to reinterpret this result from the perspective of coexistence theory. In particular, the two strategies have no niche difference, corresponding to complete niche overlap, *ρ* = 1. In contrast, the fitness difference is determined by the balance between private efficiency and the cost of cooperation:

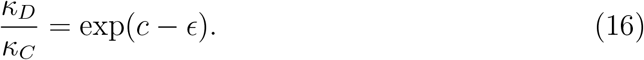

Therefore, this model always predicts competitive exclusion, except in the neutral case *c* = *ϵ*. The outcome is determined entirely by the fitness difference between cooperators and defectors, rather than by stabilizing niche differences. This example resembles the prisoner’s dilemma discussed in the previous example (Figure 3).

In real biological systems, the effect of glucose on microbial growth follows a nonlinear relationship:

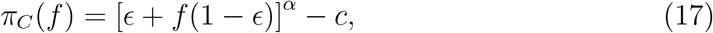

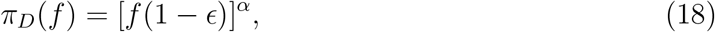

where *α* = 0.15 reflects the experimental relationship between glucose and microbial growth (Gore et al., 2009). Then, the dynamics of cooperators and defectors are described by the following replicator equation:

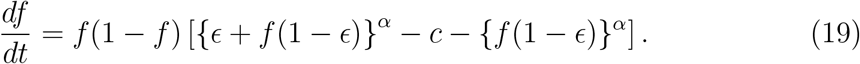

Under this nonlinear assumption, the payoff difference decreases with the cooperator fraction *f*, which can promote the coexistence of cooperators and defectors (Figure 4A): cooperators have a competitive advantage when rare, whereas defectors have a competitive advantage when cooperators are abundant.

From a coexistence-theoretic perspective, this means that nonlinear growth can generate sufficient stabilizing niche difference to overcome the fitness difference. In particular, the niche and fitness differences for this model correspond to:

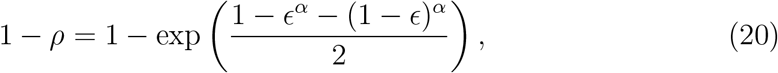

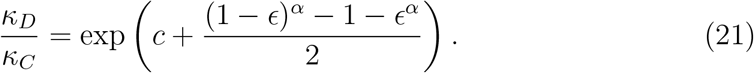

These expressions clarify the separate roles of cost, efficiency, and nonlinearity. For example, the cost of cooperation does not affect niche difference and only affects fitness difference. Instead, niche difference is generated by the concave relationship between glucose availability and microbial growth, which increases from 0 to 1 − exp(− 1*/*2) as *α* goes from 1 to 0 (Figure 4B,C). In contrast, as *α* → 0^+^, the fitness difference approaches

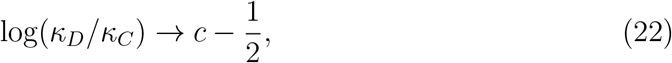

which no longer depends on the private capture efficiency *ϵ* (Figure 4C). Thus, strong concavity not only generates stabilizing niche differences, but also acts as an equalizing mechanism (Figure 4C). Such stabilizing mechanisms are key ingredients for coexistence of cooperators and defectors.

## Discussion

Standard evolutionary game theory and replicator equations provide powerful tools for predicting the outcome of strategy competition, including strategic dominance, coexistence, and bistability (Motro, 1991; Doebeli et al., 2004; Hauert and Doebeli, 2004; Doebeli and Hauert, 2005; Gore et al., 2009; Souza et al., 2009; Archetti and Scheuring, 2011; Steidinger and Bever, 2014; Tarnita and Traulsen, 2025). However, these outcomes are typically understood by analyzing payoff differences within individual games, which makes it difficult to compare coexistence mechanisms across models. Building on Chesson’s Modern Coexistence Theory from community ecology (Chesson, 2000), we developed a strategic coexistence theory (SCT) that allows us to decompose strategy competition in terms of stabilizing niche differences and fitness differences. This decomposition provides a unifying framework for comparing different games from the perspective of coexistence theory, allowing us to identify stabilizing and equalizing mechanisms for strategy competition.

To demonstrate the utility of SCT, we considered three case studies. First, the analysis of classic two-strategy games showed that SCT recovers the original classification for these classic games, highlighting the similarities between evolutionary games and community ecology (Hauert, 2002; Szabó and Fath, 2007; Cooney, 2020; Tarnita and Traulsen, 2025). Then, we compared different mechanisms for the evolution of cooperation, which showed that each mechanism promotes cooperation via different dynamical routes (Nowak, 2006). Kin selection, network reciprocity, and group selection primarily affect the fitness difference between cooperation and defection, whereas direct and indirect reciprocity can destabilize competition and generate priority effects. Finally, our analysis of microbial cooperation showed that coexistence between cooperators and defectors requires more than reducing the fitness cost of cooperation: nonlinear benefits can generate stabilizing niche differences while also reducing fitness asymmetry (Gore et al., 2009). Taken together, these examples show that SCT can provide new insights into ecological underpinnings of strategy competition.

There are several limitations to SCT. First, similar to MCT from community ecology, SCT only allows for comparisons of pairwise strategies and therefore cannot determine the outcomes of games with more than two strategies (Chesson, 2000). Second, we focused on classic replicator equations, but Tarnita and Traulsen (2025) recently pointed out that these classic formulations implicitly ignore differences in intrinsic growth rates. In such cases, replicator equations may fail to predict the outcome of competition (Tarnita and Traulsen, 2025), and a full Lotka-Volterra model is needed to describe the abundances, rather than frequencies, of strategies. Thus, SCT inherits the same assumptions as the classic replicator framework: it is most appropriate when strategy competition can be described in terms of relative frequencies alone (Tarnita and Traulsen, 2025). When strategies differ intrinsically in growth, MCT may be required instead to tease apart mechanisms of strategy coexistence. Finally, we have focused on simple examples to illustrate the utility of SCT, but many games involve additional complexities, including population structure (Nowak et al., 2010), stochasticity (Wang et al., 2025; Bodin et al., 2026), and explicit ecological feedback (Tarnita and Traulsen, 2025). Extending SCT to these settings will be necessary to determine how broadly stabilizing and equalizing mechanisms can be compared across evolutionary games.

In conclusion, SCT provides a bridge between evolutionary game theory and community ecology, allowing strategy competition to be understood from the perspective of coexistence theory. Therefore, rather than replacing existing EGT or replicator-dynamics frameworks, SCT provides a complementary tool for teasing apart the coexistence mechanisms underlying evolutionary games. More broadly, our study builds on recent efforts to extend MCT beyond species competition (Park et al., 2024, 2026), providing a foundation for building unifying coexistence theory for competing entities, including species, pathogen strains, and behavioral strategies.

## Acknowledgement

We thank Jonathan Levine for the helpful discussion. S.W.P. was supported by the New Faculty Startup Fund from Seoul National University, the National Research Foundation of Korea (NRF) grant funded by the Korea government (MSIT) (RS-2026-25474574), and the Global-LAMP Program of the National Research Foundation of Korea (NRF) grant funded by the Ministry of Education (No. RS-2023-00301976).

## Data availability

All data and code are stored in a publicly available GitHub repository (https://github.com/parklab-snu/coexistence-strategy).

## Supplementary Text

### S1 Derivation of the strategic coexistence theory

To derive strategic coexistence theory (SCT), we begin with replicator equations:

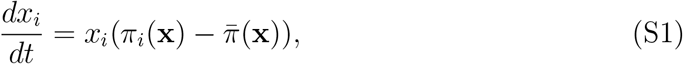

where *x*_*i*_ represents the frequency of strategy *i, π*_*i*_ represents the payoff of strategy *i*, and 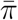 represents the average payoff. Then, the mutual invasion condition for two competing strategies *i* and *j* can be written as

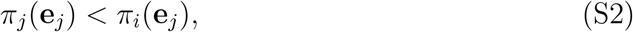

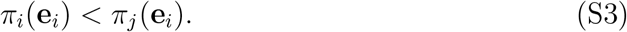

These inequalities will be preserved when we exponentiate the payoffs, *W*_*i*_(**x**) = exp(*π*_*i*_(**x**)):

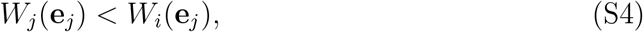

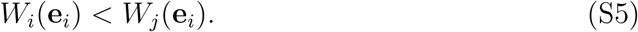

Rearranging, we obtain the following inequality:

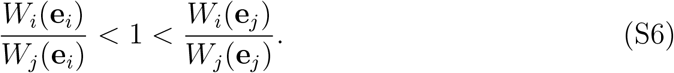

By multiplying 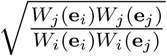 on both sides, we obtain:

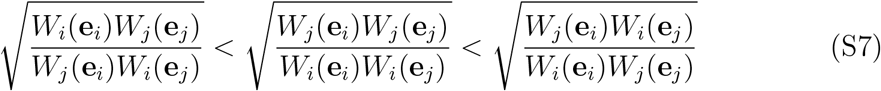

Thus, by defining niche overlap *ρ* and fitness difference *κ*_*j*_*/κ*_*i*_ as follows,

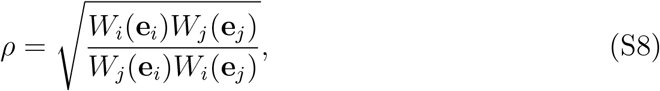

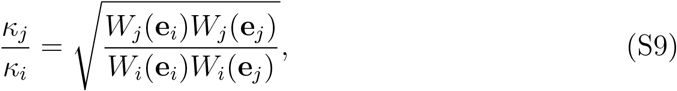

we obtain the following inequality that defines conditions for mutual invasion and long-term coexistence of two competing strategies:

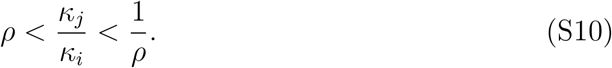

### S2 Applications to classic two-strategy games

Consider a two-strategy game described by the following payoff matrix:

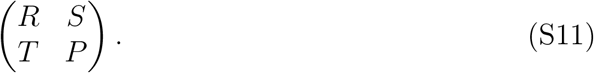

There are four classic games that we can define from this simple payoff matrix: prisoner’s dilemma (*T > R > P > S*), hawk–dove game (*T > R, S > P* ), stag–hunt game (*R > T, P > S*), and harmony game (*R > T, S > P* ). Here, we show that each game corresponds to a distinct region in the niche-fitness difference space under SCT.

To do so, we first begin by deriving expressions for niche overlap *ρ* and fitness difference *κ*_*j*_*/κ*_*i*_. Writing *x*_1_ to denote the frequency of strategy 1, the expected payoff for each strategy corresponds to:

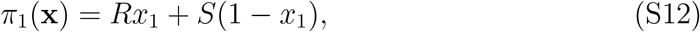

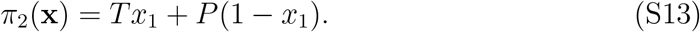

Then, multiplicative payoffs for each strategy when rare can be written as:

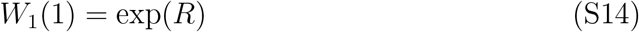

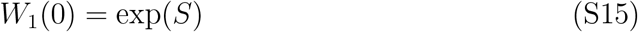

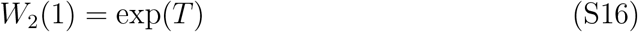

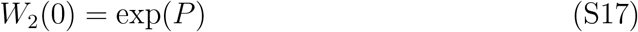

Therefore, we have:

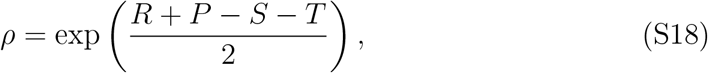

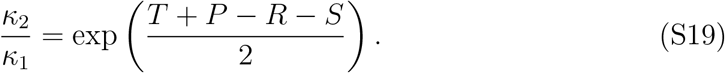

As before, SCT allows us to predict the mutual invasion of two strategy using the following inequality:

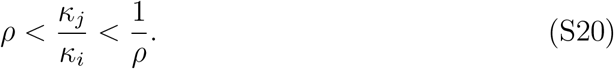

Then, simple algebra reveals that each condition in mutual invasion inequality can be equivalently described by the payoff inequality:

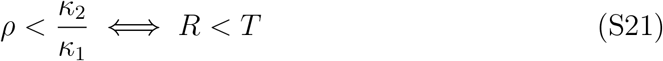

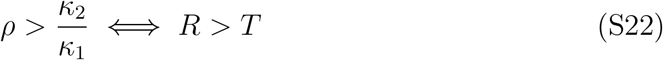

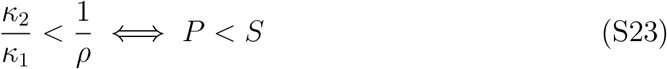

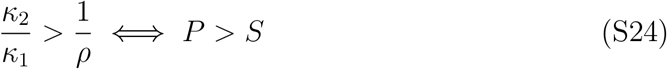

These inequalities then allow us to classify outcomes of two-strategy games from the perspective of SCT. Note that the inequality between *R* and *P* does not affect the outcome.

### S3 Applications to mechanisms for the evolution of cooperation

We consider five mechanisms for the evolution of cooperation, all of which can be represented as a simple 2 × 2 payoff matrix (Nowak, 2006). Here, we go through each mechanism and derive measures for niche and fitness differences.

#### Prisoner’s dilemma

Here, we consider a special case of Prisoner’s dilemma which can be described by the following payoff matrix:

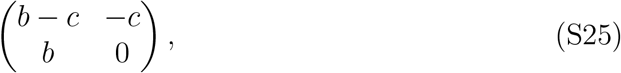

where *b* represents the benefit for the recipient and *c* represents cost for the donor. In this case, the niche and fitness differences correspond to:

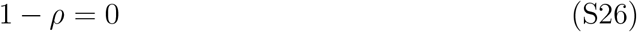

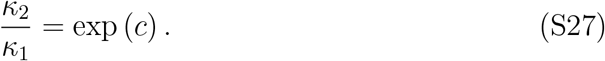

Therefore, as long as the cost is greater than zero, prisoner’s dilemma always results in the competitive exclusion of cooperation.

#### Kin selection

Kin selection can be described by the following payoff matrix:

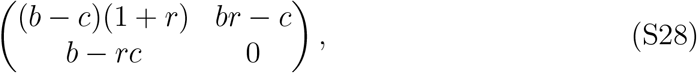

where *r* represents the genetic relatedness. In this case, the niche and fitness differences correspond to:

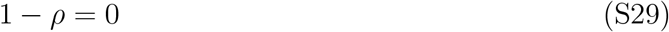

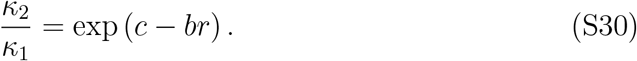

Therefore, defection will exclude cooperation when *c > br*. Equivalently, cooperation will be favored when *r > c/b*, which recovers the classic result.

#### Direct reciprocity

Direct reciprocity can be described by the following payoff matrix:

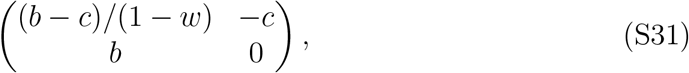

where *w* represents the probability of next round. In this case, the niche and fitness differences correspond to:

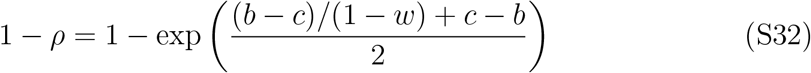

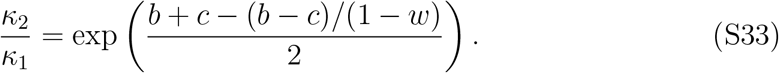

Therefore, defection will exclude cooperation when

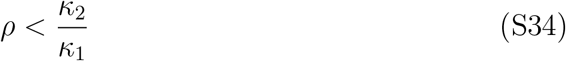

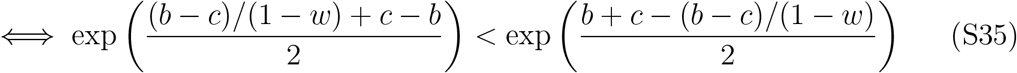

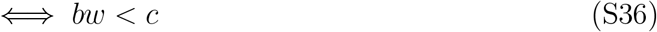

However, note that the following equality holds whenever *c >* 0:

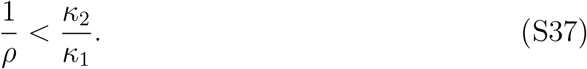

Therefore, when *w > c/b*, we will have

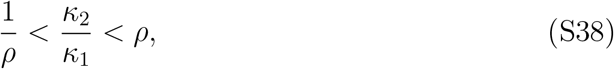

which implies priority effect (equivalently, bistability). Even as *w* → 1^*−*^, this inequality holds and therefore, cooperation cannot competitively exclude defection.

#### Indirect reciprocity

Indirect reciprocity can be described by the following payoff matrix:

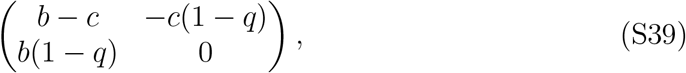

where *q* represents social acquaintanceship. In this case, the niche and fitness differences correspond to:

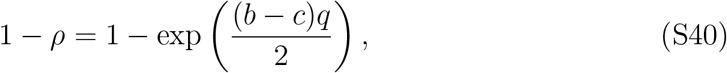

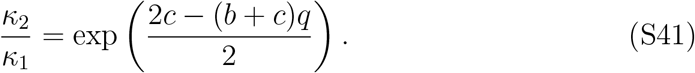

In this case, the condition for the dominance of defection (and therefore competitive exclusion of cooperation) can be written as:

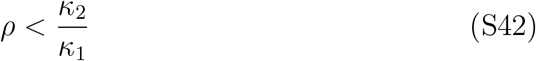

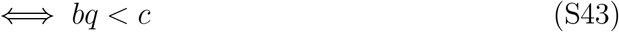

Moreover, the following inequality always holds as long as *c >* 0.

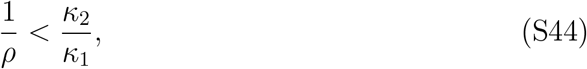

which implies that coexistence between cooperation and defection is not possible. Therefore, when *q < c/b*, then defection will competitively exclude cooperation. Otherwise, the system will enter the priority-effect regime.

Interestingly, when *q* → 1, we have

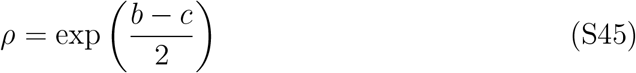

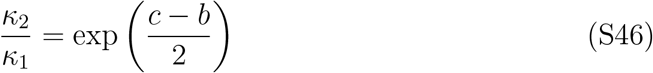

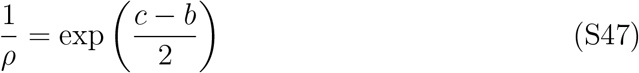

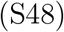

In other words, the following inequality will hold:

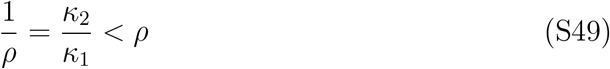

#### Network reciprocity

Network reciprocity can be described by the following payoff matrix:

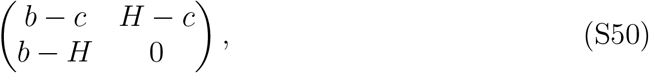

where *H* = [(*b* − *c*)*k* − 2*c*]*/*[(*k* + 1)(*k* − 2)] and *k* represents the number of neighbors. In this case, the niche and fitness differences correspond to:

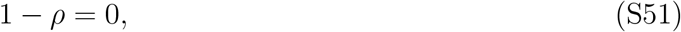

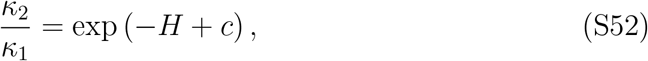

which implies that there is complete niche overlap and that fitness difference decreases with *H*. Since *H* is a monotonically decreasing function of *k* for *k >* 2, fitness difference will increase with *k*. In particular, cooperation will competitively exclude defection when *H > c*, meaning that:

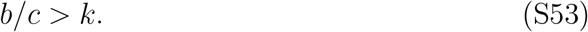

#### Group selection

Group selection can be described by the following payoff matrix:

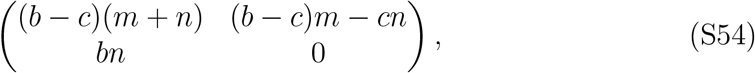

where *m* represents the number of groups and *n* represents the group size. In this case, the niche and fitness differences correspond to:

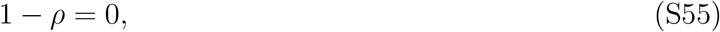

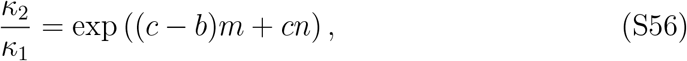

which implies that there is complete niche overlap. The fitness difference will increase with *n* but decrease with *m*, provided that *b > c*. In particular, cooperation will competitively exclude defection under the following condition:

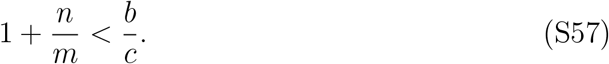

